# The infant airway microbiome in health and disease impacts later asthma development

**DOI:** 10.1101/012070

**Authors:** Shu Mei Teo, Danny Mok, Kym Pham, Merci Kusel, Michael Serralha, Niamh Troy, Barbara J. Holt, Belinda J. Hales, Michael L. Walker, Elysia Hollams, Yury H Bochkov, Kristine Grindle, Sebastian L. Johnston, James E Gern, Peter D. Sly, Patrick G. Holt, Kathryn E. Holt, Michael Inouye

## Abstract

The nasopharynx (NP) is a reservoir for microbes associated with acute respiratory illnesses (ARI). The development of asthma is initiated during infancy, driven by airway inflammation associated with infections. Here, we report viral and bacterial community profiling of NP aspirates across a birth cohort, capturing all lower respiratory illnesses during their first year. Most infants were initially colonized with *Staphylococcus* or *Corynebacterium* before stable colonization with *Alloiococcus* or *Moraxella*, with transient incursions of *Streptococcus, Moraxella* or *Haemophilus* marking virus-associated ARIs. Our data identify the NP microbiome as a determinant for infection spread to the lower airways, severity of accompanying inflammatory symptoms, and risk for future asthma development. Early asymptomatic colonization with *Streptococcus* was a strong asthma predictor, and antibiotic usage disrupted asymptomatic colonization patterns.

## Introduction

The microbial communities comprising the human microbiome are now recognised as playing important roles in the aetiology and pathogenesis of myriad diseases (*1*). However elucidation of these complex roles requires targeted characterization of microbial communities present in relevant spatial niche(s) during critical periods of pathogenesis. The focus of this study is the respiratory tract, in particular the nasopharynx (NP) which is an accessible source of airway microbial communities (*2*) and serves as a conduit for pathogens associated with lower respiratory illnesses (LRI) that are responsible for substantial morbidity and mortality worldwide.

Of particular interest is asthma, a multi-factorial disease characterized by airway inflammation and associated smooth muscle hyperplasia. It is now recognized that the hallmark persistent wheeze of asthma is consolidated in childhood and, further, may progress to chronic asthma in adulthood (*3*,*4*) and potentially chronic obstructive pulmonary disease (*5*, *6*). We and others have previously shown that development of persistent atopic (allergic) wheeze in children is linked to the number of virus-associated febrile and/or wheezy LRI experienced during infancy (*7*-*10*). The principal virus type of current interest is human rhinoviruses (HRV), particularly subtype C (HRV-C) (*11*), however respiratory syncytial virus (RSV) is also recognized as a major cause of infant LRI (*12*). The relative contributions of these viral pathogens in asthma initiation remain controversial (*13*). Further complicating the picture, recent studies have also implicated bacterial pathogens as potential independent causal factors in infant LRI and their long-term sequelae. Notably, culture of *S. pneumoniae, M. catarrhalis* or *H. influenzae* from NP samples taken at 1 month of age has been linked to increased risk for subsequent diagnosis of asthma at 5 years of age (*14*). These findings have fuelled debate around the use of antibiotics and vaccine strategies for respiratory illness in children (*15*, *16*).

Several studies have investigated airway microbiota in children or adults with chronic respiratory illness, including asthma (*2*, *17*-*21*), however no study has investigated the airway microbiome during the critical infancy period (0-12 months). In this study, we investigated the NP microbiome during the first year of life using the Childhood Asthma Study (CAS), a prospective cohort of 234 children (*9*, *10*, *22*, *23*), to elucidate the NP microbiome during respiratory health and illness, its longitudinal dynamics, susceptibility to exogenous factors such as antibiotics, and association with future asthma.

## Results

A total of 1,021 NP microbiome profiles were obtained from 234 infants using 16S rRNA gene deep sequencing (**Materials and Methods**). These included 487 “healthy” NP samples collected in the absence of respiratory symptoms, and 534 “infection” NP samples collected during episodes of acute respiratory illness (ARI) during the first year of life. Three quarters of the infants (N=177) contributed at least two healthy NP samples at the age of ~2 months, ~6 months and/or ~12 months. Eighty percent of the infants (N=186) contributed a healthy sample before experiencing their first ARI. The 534 infection NP samples were from 184 infants who had experienced ≥1 ARI within the first year of life. NP samples were analysed from all (380/381) recorded LRI in this period, and a random selection of 20% (154/782) of recorded episodes of upper respiratory illness (URI). The characteristics of the infants are summarized in **Table S1**.

### NP microbiome composition

Across all NP samples, more than 193 million high quality 16S rRNA sequences were classified into 14,131 operational taxonomic units (OTUs), of which 1,010 were supported by >1,000 reads each. The dominant phyla were Proteobacteria (48%), Firmicutes (38%), Actinobacteria (13%), Bacteroidetes (1%) and Fusobacteria (0.5%) (**Fig. 1A**). The NP microbiomes were dominated by six genera - *Moraxella* (31.2%), *Streptococcus* (15.5%), *Corynebacterium* (13.5%), *Staphylococcus* (10.3%), *Haemophilus* (9.7%) and *Alloiococcus* (8.8%; grouped under genus *Dolosigranulum* in some databases) (Error! Reference source not found.**A**). Despite the inclusion of diverse OTUs of these genera in the Greengenes reference database (138 *Moraxella*; 2,361 *Streptococcus*; 1,592 *Corynebacterium*; 1,936 *Staphylococcus*; 684 *Haemophilus*; 82 *Alloiococcus*), our sequences were dominated by one OTU per genus (**Fig. S1-6**), consistent with culture-based studies reporting NP colonization with the species *Moraxella catarrhalis*, *Streptococcus pneumoniae, Staphylococcus aureus*, *Haemophilus influenzae* and *Alloiococcus otitidis*. Hierarchical clustering of NP microbiomes based on relative abundance of the six major genera identified six microbiome profile groups (MPGs, **Fig. 1B**). Each MPG was dominated by one of the six genera, although some samples in the *Alloiococcus* MPG also had relatively high abundance of *Corynebacterium* (**Fig. 1B**). The distribution of other rarer OTUs, including those of *Neisseria*, *Pseudomonas*, *Veillonella*, *Gemella*, *Rothia* and *Prevotella*, is shown in **Fig. S7**.

**Fig. 1.**
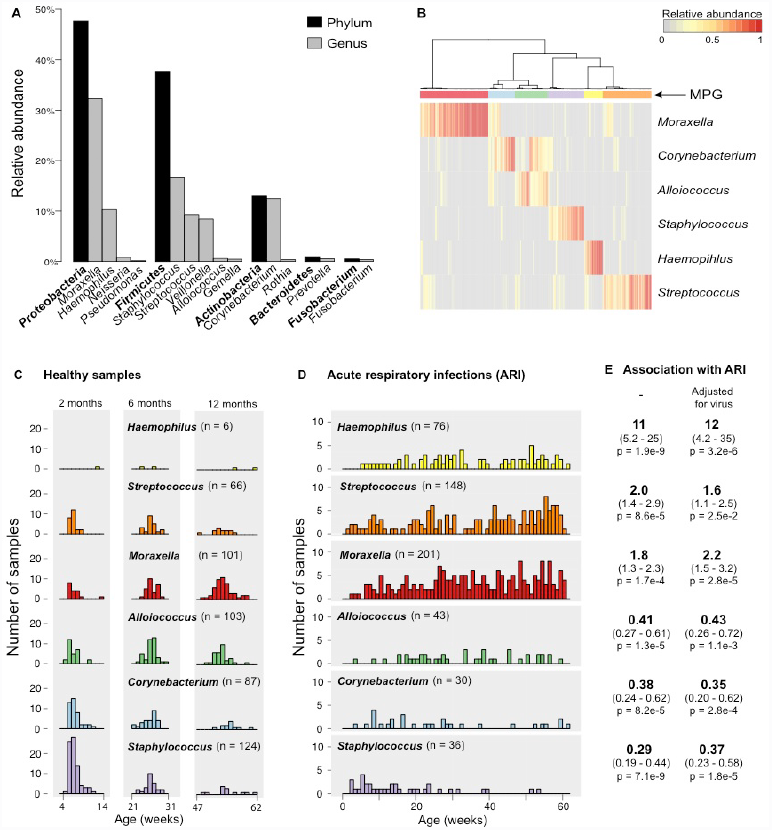
Bacterial composition of 1,021 nasopharyngeal aspirates collected from 234 infants during periods of respiratory health and disease. **(A)** Frequency of the 5 most abundant phyla and their respective genera (taxa shown comprise 99.9% of all reads). **(B)** Hierarchical clustering of samples into microbiome profile groups (MPGs) based on relative abundance of the 6 most common genera. Coloured bars indicate MPGs, labelled throughout the text by their dominant genus: *Moraxella* MPG (red), *Corynebacterium* MPG (blue), *Alloiococcus* MPG (green), *Staphylococcus* MPG (purple), *Haemophilus* MPG (yellow) and *Streptococcus* MPG (orange). **(C)** Weekly frequencies of MPGs amongst healthy samples, collected during planned visits at approximately 2 months, 6 months and 12 months of age and following at least 4 weeks without symptoms of acute respiratory infection (ARI). **(D)** Weekly frequencies of each MPG amongst ARI samples. **(E)** Odds ratios for association of MPGs with ARI symptoms (i.e., ARI vs healthy samples), adjusted for age, gender, season, number of prior infections, antibiotics intake, mother’s antibiotics intake, delivery mode and breastfeeding; with and without adjustment for detection of common viruses (RSV, HRV).

### NP microbiome dynamics

Healthy NP samples collected around 2 months of age were dominated by the *Staphylococcus* (41%) and *Corynebacterium* (22%) MPGs, but the frequency of these MPGs declined with age (11% and 10%, respectively, at 12 months old) (**Fig. 1C**). In contrast, the prevalence of *Alloiococcus* and *Moraxella* MPGs in healthy samples increased with age (14% and 9% at 2 months, 26% and 41% at 12 months, respectively). Analysis of MPG transitions amongst consecutive healthy NP samples from the same individuals (**Fig. S8A-B**) suggested that *Staphylococcus* carriage was unstable or transient, particularly where an ARI had occurred in the intervening period between sampling. *Alloiococcus* was a stable colonizer, but less so if an ARI occurred between sampling. Where an ARI occurred in the intervening period between healthy samples, the most common transitions were to the *Moraxella* MPG from other MPGs, or the maintenance of stable colonization with *Moraxella* (**Fig. S8B**). Almost all *Moraxella*-colonized infants experienced subsequent ARI before the next healthy sample was taken; this is explored further below. *Haemophilus* was very rarely detected in healthy NP microbiomes, while *Streptococcus* was present in 14% of healthy samples at each of the 2, 6 and 12 month sampling times (**Fig. 1C**). Similar age-related patterns were observed amongst infection samples, with a decline in *Staphylococcus* and *Corynebacterium* MPGs and increase in *Moraxella*, *Haemophilus* and *Streptococcus* MPGs in older children (**Fig. 1D**). Interestingly, boys had significantly more *Moraxella* in healthy samples (OR 1.3 for log abundance; 95% CI 1.1-1.5, p=0.0014, adjusted for age); no other gender effects were detected.

We next assessed the impact of environmental factors on the relative abundance of the common NP microbiome genera. We found no significant effects of delivery mode or breastfeeding on NP colonization at 2 months of age; the latter was unsurprising as nearly all infants (90%) were breastfed for at least 2 months. The abundance of *Streptococcus* in healthy NP samples was significantly lower amongst children whose parents reported having furry pets such as dogs or cats in the home (OR 0.84, 95% CI 0.70-1.0, p=0.046; adjusted for age at NP sampling). No other significant associations were detected for pets. Children attending day-care had significantly higher relative abundances of *Haemophilus* and *Moraxella*, and lower relative abundances of *Corynebacterium* and *Staphylococcus* (**Fig. 2A**), in both healthy and infection samples (note that very few children had commenced day-care by 6 months of age, hence the impact of day-care attendance was assessed at the 12-month time point only). Co-habiting with siblings was also associated with higher abundances of *Haemophilus*, *Streptococcus* and *Moraxella*, and lower abundance of *Staphylococcus*, during health and ARI (adjusted for age at sampling, **Fig. 2B**). Importantly, amongst healthy samples, antibiotic usage in the four weeks prior to sampling was associated with higher abundances of *Haemophilus, Streptococcus* and *Moraxella*, and lower abundances of *Alloiococcus* and *Corynebacterium* (adjusted for age at sampling, **Fig. 2C**). The composition of the healthy NP microbiome was also affected by the number of prior respiratory infections experienced, with higher abundance of *Moraxella* and lower abundances of *Alloiococcus* or *Corynebacterium* in samples following increasing numbers of ARIs (**Fig. S9**). At the MPG level, ARIs dominated by *Haemophilus* increased in spring-summer, while those dominated by *Moraxella* peaked in autumn-winter (**Fig. S10**). We therefore tested for differences in relative abundance in autumn-winter and spring-summer, adjusting for age and number of prior infections; this confirmed significant seasonal effects on the abundance of *Haemophilus* (summer associated) and *Moraxella* (winter associated) amongst ARI samples, and a similar but non-significant trend amongst healthy NP samples (**Fig. 2D**).

**Fig. 2.**
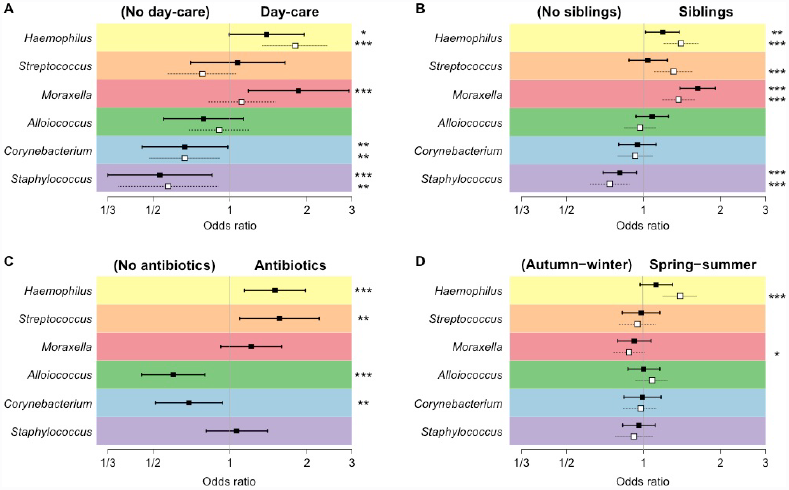
Impact of environmental factors on relative abundances of major genera in the NP microbiome. Squares, odds ratios estimated using logistic regression and adjusted for age; filled squares, healthy samples; empty squares, infection samples; bars, 95% confidence intervals; *p<0.1, **p<0.05, ***p<0.01. **(A)** Day-care attendance (yes vs. no), amongst 12-month samples only. **(B)** Co-habiting with siblings (yes vs. no). **(C)** Antibiotics intake in the 4 weeks preceding NP sample collection (yes vs. no). **(D)** Season; Spring-summer (September-February) vs. Autumn-winter (March-August).

### NP microbial determinants of ARI symptoms

The *Moraxella*, *Streptococcus* and *Haemophilus* MPGs were significantly more frequent in ARI compared to healthy NP samples, even after adjusting for a large set of potential confounders (age, gender, season, number of prior infections, antibiotic intake, mother’s antibiotic intake, delivery mode and breastfeeding) (**Fig. 1E**). The *Staphylococcus, Corynebacterium* and *Alloiococcus* MPGs were significantly less frequent in ARI (**Fig. 1E**). The rare genus *Neisseria* was more common in infection samples, especially LRIs (of 28 NP samples with >5% relative abundance of *Neisseria*, 4 were from URI and 20 were from LRI), though all were co-colonized with *Streptococcus*). It was not possible to confirm species from the 16S sequences, however it is well-known that the respiratory pathogens *S. pneumoniae*, *H. influenzae*, *M. catarrhalis* and *N. meningitidis* are frequently cultured from respiratory infections in children. We have previously measured IgG to species-specific surface proteins of *S. pneumoniae* and *H. influenzae* at 12 months of age in the CAS cohort (*24*). Here we found *H. influenzae*-specific IgG was significantly associated with the number of prior ARI samples testing positive for either of the two most common *Haemophilus* OTUs (**Fig. S1**); similar results were obtained for *S. pneumoniae*-specific IgG antibodies and the dominant *Streptococcus* OTU (**Fig. S3**). Healthy colonization with these genera was not associated with species-specific IgG.

A total of 138 children had ≥2 ARI samples profiled and, of these, 97 individuals (70%) had ≥2 different MPGs amongst their ARIs. There was a clear temporal trend with infections occurring closer in time being more likely to cluster into the same MPG (same MPG in half of infection pairs separated by 5 weeks vs. a quarter of those separated by >20 weeks; **Fig. S11**). *Moraxella* and *Haemophilus* MPGs were particularly stable between consecutive infections, i.e. following an ARI in which the *Moraxella* or *Haemophilus* MPG was present, the next ARI was more likely to share the same MPG type than expected given the overall frequency of these MPGs amongst infection samples (**Fig. S8C**).

We further considered the impact of the NP microbiome on infection severity and interactions with viral pathogens. All samples analysed here were previously screened for a panel of viruses, including RSV, HRV and other picornaviruses, influenza, parainfluenza, coronavirus, adenovirus and human metapneumovirus (hMPV) (see **Materials and Methods** (*23*)). This first screen detected viruses in 21% of healthy samples, 68% of URI and 69% of LRI. The most common viruses detected were RSV (11% of ARI) and HRV (40% of ARI) (**Fig. S12B**), although subsequent expanded screening and subtyping of HRV in LRI samples suggests this is an underestimate (see below). For all virus groups except adenovirus and coronavirus, virus detection was significantly positively associated with ARI symptoms (i.e. ARI vs healthy samples, **Fig. S12B**). The association between ARI and *Streptococcus*, *Haemophilus* and *Moraxella* MPGs remained after adjusting for detection of virus (OR 7.0, p<1x10^-15^ for any of these MPGs; individual ORs in **Fig. 1E**), indicating both viruses and bacteria contribute to ARI symptoms. Amongst the viruses analysed, only RSV was significantly more frequent in LRI vs URI (16% vs 8.3%, OR 2.3; **Fig. S12B**, **Table 1**). The *Streptococcus*, *Haemophilus* and *Moraxella* MPGs were significantly associated with LRI vs URI (OR >2), individually and collectively, adjusting for the effect of RSV (**Table 1**). As a group, the illness-associated MPGs *Streptococcus*, *Haemophilus* and *Moraxella* increased in frequency from healthy to URI to LRI samples, regardless of the presence of RSV (**Fig. S13**). Taken together, these analyses indicate that both viruses and bacteria independently contribute to ARI, and that bacteria and RSV also independently increase the risk of infection spread to the lower airways.

**Table 1.** Associations between ARI symptom severity and risk-associated bacterial MPGs. Odds ratio (OR) (95% confidence interval), p-values. NP samples taken within one week of antibiotic use were excluded from analysis. The response variable (symptom group comparison) is shown in column 1. Two models (A, B) were fit for each comparison, labelled in column 2; Model A included *Streptococcus* or *Moraxella* or *Haemophilus* MPG as a single covariate, and RSV and HRV, further adjusting for age, gender and season; Model B included *Streptococcus*, *Moraxella* and *Haemophilus* MPGs as separate covariates, and RSV and HRV, further adjusting for age, gender and season. Note HRV was not included in the LRI vs URI comparison as enhanced sensitivity re-screening for HRV was performed only in LRI samples.

| Symptom group comparison | Model | *Streptococcus, Moraxella or Haemophilus* MPG |  |  | RSV | HRV (expanded screen) |
| --- | --- | --- | --- | --- | --- | --- |
|  |  | *Streptococcus* MPG | *Moraxella* MPG | *Haemophilus* MPG |  |  |
| **LRI vs. URI** | 1A | OR = 2.2 (1.3 - 3.7), p = 0.0043 |  |  | 2.2<br>(1.1 - 4.7)<br>p = 0.034 | - |
|  | 1B | 2.3<br>(1.2 - 4.5)<br>p = 0.011 | 2.1<br>(1.1 - 3.8)<br>p = 0.017 | 2.1<br>(1.0 - 4.6)<br>p = 0.056 | 2.3<br>(1.1 - 4.8)<br>p = 0.033 | - |
| **Febrile LRI vs. non-febrile LRI** | 2A | OR = 3.8 (1.0-14), p = 0.047 |  |  | 4.0<br>(1.6 - 9.8)<br>p = 0.0031 | 0.45<br>(0.21 - 0.95)<br>p = 0.036 |
|  | 2B | 4.3<br>(1.0 - 18)<br>p = 0.046 | 3.8<br>(0.98 - 15)<br>p = 0.053 | 2.7<br>(0.53-14)<br>p = 0.23 | 4.0<br>(1.6 - 10)<br>p = 0.0031 | 0.46<br>(0.21 - 0.98)<br>p = 0.043 |
| **Wheezy LRI vs. non-wheezy LRI** | 3A | OR = 1.4 (0.60-3.1), p = 0.45 |  |  | 0.94<br>(0.41 - 2.1)<br>p = 0.88 | 1.2<br>(0.62 - 2.2)<br>p = 0.64 |
|  | 3B | 1.1<br>(0.39 - 3.0)<br>p = 0.88 | 1.8<br>(0.74 - 4.1)<br>p = 0.20 | 1.0<br>(0.34 - 3.2)<br>p = 0.93 | 0.95<br>(0.42 - 2.2)<br>p = 0.90 | 1.1<br>(0.60 - 2.2)<br>p = 0.69 |

In our earlier studies with this cohort, we observed an association between LRI (but not URI) during infancy and risk for wheeze at age 5 years; moreover this association was restricted to severe LRI, i.e. those accompanied by fever and/or wheeze (*9*,*10*,*25*). Thus we investigated the association of viruses and bacteria with the presence of fever and wheeze symptoms during LRI. To enable more accurate assessment of the role of HRV in this critical sample group, we performed an expanded screen for detection and subtyping of HRV within LRI samples (**Materials and Methods**). This assay proved more sensitive than the first screen (*23*) and detected HRV in 66% of LRI samples (compared to 33% in the earlier screen), with equal amounts of HRV-A and HRV-C (**Fig. S12**). During LRI, the presence of any HRV, or HRV-C specifically, was negatively associated with fever (**Table 1**). The only viruses showing positive associations with fever were RSV (**Table 1**, **Fig. 3C**, **Fig. S12**) and influenza (8 influenza positive fever LRI events only, all with illness-associated MPGs, thus not considered further). Amongst LRI, the illness-associated MPGs were associated with fever, even after adjusting for RSV, HRV, age, season and gender (**Table 1**). Interestingly, the *Moraxella* MPG was also significantly positively associated with fever amongst RSV-positive LRI (OR 9.2, 95% CI 1.1-73, p=0.037; **Fig. S13**), suggesting a possible interaction between *Moraxella* and RSV whereby their co-presence further enhances risk of fever. Overall, only 10 febrile LRIs (9.5%) could not be explained by the presence of RSV or illness-associated MPGs. Across the cohort, the presence of wheeze during LRI was not significantly associated with any viral or bacterial groups (**Table 1**), including HRV or HRV-C specifically. Of all ARI analysed, there were 61 with concomitant otitis media (OM) affecting 47 infants, however ARIs with accompanying OM had similar NP microbiome profiles to those of LRIs with no OM diagnosis (**Table S2**).

**Fig. 3.**
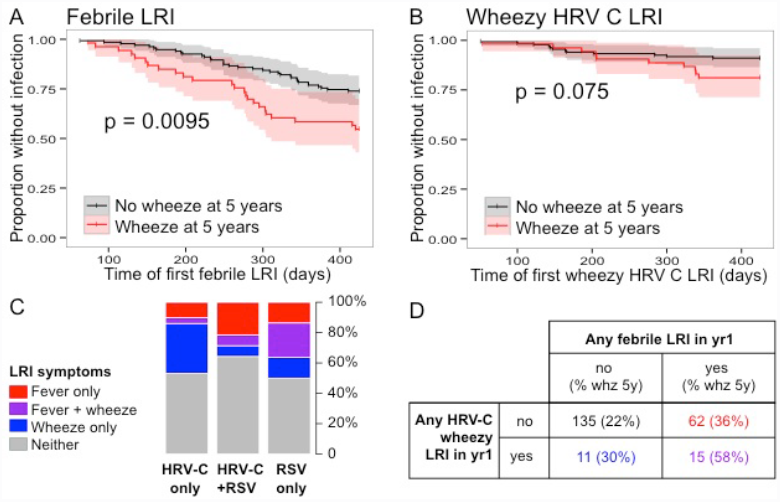
Symptoms of lower respiratory illness (LRI) during the first year of life are associated both with viruses present during the infection, and with chronic wheeze at 5 years of age. **(A-B)** Kaplan-Meier survival curves showing time from birth to **(A)** first febrile LRI and **(B)** first HRV-C wheezy LRI, stratified according to the child’s chronic wheeze status at 5 years of age. P-values shown were estimated using Cox proportional hazards models, adjusted for gender and maternal and paternal history of atopic disease. **(C)** Frequencies of fever and wheeze symptoms during LRI, in which HRV-C and/or RSV were detected. Total numbers are: HRV-C only, n=79; HRV-C + RSV, n=14; RSV only, n=22. **(D)** Cross-tabulation of individuals, stratified by their experience of LRI during infancy; percentages in brackets indicate the frequency of chronic wheeze at 5 years of age within each group.

### Impact of ARI on later chronic wheeze

We next investigated associations between LRI in the first year of life and subsequent expression of chronic wheeze at age 5 or 10 years. Positive associations were found for two discrete classes of LRI. Firstly, febrile LRI was significantly positively associated with later chronic wheeze, and amongst children who developed atopy by 2 years of age (**Table 2**). Furthermore, timing of first febrile LRI appeared to be important, with earlier febrile LRI occuring amongst children who had chronic wheeze at 5 years (p=0.0095, **Fig. 3A**). Secondly, HRV-C LRI accompanied by wheezing symptoms (wheezy HRV-C LRI) showed a positive association with later chronic wheeze amongst all children and particularly strongly so for those who were atopic by 2 years (OR ~7, **Table 2**), but not amongst non-atopics (OR 1, p~0.95; **Table S3**). As we found no association between HRV-C and wheeze during LRI across the whole cohort (OR 1.2, 95% CI 0.63-2.2, p=0.61; adjusting for illness-associated MPGs, RSV, age, season and gender), we re-examined the association in those who developed atopy by age 2. Amongst this group of children (n~65), the presence of HRV-C was significantly positively associated with increased risk of wheeze during LRI (OR 2.7, 95% CI 1.1-7, p~0.035; adjusting for illness-associated MPGs, RSV, age, season and gender).

**Table 2.** Association between microbial events during infancy and chronic wheeze at age 5 and 10 years. Odds ratios, 95% confidence intervals and p-values shown were estimated using logistic regression, adjusted for gender and maternal and paternal history of atopic disease, estimated separately for all children and those who were atopic by 2 years of age. LRI, lower respiratory illness, further classified according to microbes and symptoms; any risk bacteria LRI, any LRI with *Streptococcus*, *Moraxella* or *Haemophilus* MPG. Early *Streptococcus* colonization was assessed in the first healthy NP sample, collected by 7 weeks of age and prior to any recorded infection; high abundance was classified as >20% *Streptococcus* reads (based on distribution shown in **Fig. S15**).

|  | Wheeze at 5 years |  | Wheeze at 10 years |  |
| --- | --- | --- | --- | --- |
|  | All children | Atopics | All children | Atopics |
| **Any febrile LRI** | 2.3<br>(1.2 - 4.5)<br>p = 0.016* | 2.7<br>(1.1 - 7)<br>p = 0.034* | 2.2<br>(0.93 - 5.2)<br>p = 0.071 | 2.7<br>(0.89 - 8.7)<br>p = 0.083 |
| **Any wheezy LRI** | 1.6<br>(0.79 - 3)<br>p = 0.2 | 1.3<br>(0.49 - 3.3)<br>p = 0.61 | 1.4<br>(0.58 - 3.3)<br>p = 0.45 | 1.6<br>(0.53 - 4.8)<br>p = 0.4 |
| **Any HRV wheezy LRI** | 2<br>(0.93 - 4.2)<br>p = 0.073 | 2.5<br>(0.86 - 7.2)<br>p = 0.092 | 2.1<br>(0.81 - 5.4)<br>p = 0.11 | 1.9<br>(0.55 - 6.4)<br>p = 0.29 |
| **Any HRV-C wheezy LRI** | 2.4<br>(0.93 - 6.1)<br>p = 0.064 | 7.2<br>(1.7 - 35)<br>p = 0.009* | 3.5<br>(1.1 - 11)<br>p = 0.026* | 7.1<br>(1.6 - 40)<br>p = 0.014* |
| **Any HRV-A wheezy LRI** | 1.2<br>(0.42 - 3.1)<br>p = 0.74 | 0.55<br>(0.11 - 2.2)<br>p = 0.43 | 1.4<br>(0.37 - 4.7)<br>p = 0.57 | 0.35<br>(0.02 - 2.2)<br>p = 0.34 |
| **Any risk bacteria LRI** | 0.89<br>(0.45 - 1.8)<br>p = 0.73 | 0.96<br>(0.39 - 2.4)<br>p = 0.93 | 2<br>(0.82 - 5.2)<br>p = 0.14 | 1.7<br>(0.59 - 5.2)<br>p = 0.34 |
| **High abundance**<br>***Streptococcus* colonization**<br>**(≤7 weeks)** | 3.8<br>(1.3 - 12)<br>p = 0.017* | 4<br>(0.88 - 21)<br>p = 0.077 | 2.7<br>(0.6 - 12)<br>p = 0.18 | 3.9<br>(0.63 - 28)<br>p = 0.15 |

Since febrile LRI and HRV-C wheezy LRI were both associated with later chronic wheeze (**Table 2**) and RSV was associated with febrile LRI (**Table 1**), we examined the interaction between RSV, HRV-C, LRI symptom severity and chronic wheeze at age 5. Febrile LRI and wheezy HRV-C LRI in the first year of life appeared to exert independent effects on later chronic wheeze, since only a minority of HRV-C wheezy infections were also febrile (**Fig. 3C**). Further, at the level of individual children, the frequency of chronic wheeze at 5 years was elevated amongst those with either one of febrile LRI (36%) or wheezy HRV-C LRI (30%), but was greatest amongst children who experienced both (58%, **Fig. 3D**).

### Impact of NP colonization on ARI

Where *Streptococcus*, *Haemophilus* or *Moraxella* MPGs were detected in healthy NP samples, subsequent infections tended to belong to the same MPG (**Fig. S14A**). Critically, we investigated whether healthy colonization in early infancy was associated with subsequent episodes of ARI. Since prior infections also affect the healthy NP microbiome (**Fig. S9**), we restricted these analyses to healthy samples collected at 5-9 weeks of age and prior to each infant’s first reported ARI (n=160). Using Cox proportional hazards models, infants whose earliest healthy NP sample was of the *Moraxella* or *Streptococcus* MPG tended to experience ARI at a younger age than those with other MPGs (p=0.052; **Fig. S14B**; note *Haemophilus* MPGs were extremely rare in early healthy samples). When examining URIs and LRIs separately, we found that early *Moraxella* colonization was moderately associated with earlier first URI (p=0.084), while early *Streptococcus* colonization was strongly associated with earlier first LRI (p=0.0071; **Fig. S14C-D**). Since our data suggested that *Alloiococcus* and *Moraxella* were key stable colonizers of the NP microbiome, we also examined whether *Alloiococcus*-colonized infants differed from *Moraxella*-colonized infants (defined as those with ≥1 healthy sample of *Moraxella* MPG and none with *Alloiococcus* MPG, and vice versa). These groups did not differ in terms of overall numbers of ARI, LRI or later wheezing phenotypes. However, compared to either *Moraxella*-colonized infants or those not in either group, *Alloiococcus*-colonized infants had fewer RSV infections, especially RSV LRIs (OR 0.27, 95% CI 0.078-0.73, p=0.0050) (**Table S4**).

### Early NP colonization impacts later chronic wheeze

We next assessed the association between early pre-ARI asymptomatic NP colonization and current wheeze at 5 and 10 years of age, stratified by atopic sensitization by 2 years of age (see **Materials and Methods**). This analysis was restricted to the 160 infants (70%) who had an asymptomatic NP sample taken prior to their first ARI (and ≤9 weeks of age). *Haemophilus* was not detected in pre-ARI healthy samples and the *Moraxella* MPG showed no evidence of association with future wheeze. The *Streptococcus* MPG showed a weak association, however the relative abundance of *Streptococcus* in these samples was highly skewed, thus we divided samples into those with high (>20%) or low (≤20%) relative abundance of *Streptococcus* reads (**Fig. S15**). High *Streptococcus* abundance in the first pre-ARI healthy NP sample was more frequent in infants who later displayed chronic wheeze at 5 years, and this association was stronger when restricting the analysis to earlier NP samples (**Fig. 4A**): OR 3.8 amongst healthy NP samples collected ≤7 weeks post birth (**Table 2**). The same trend was evident for wheeze at 10 years of age, despite reduced sample size due to loss to follow-up (OR 2.7, **Table 2**). Early *Streptococcus* colonization was associated with younger age at first LRI (**Fig. S14D**), but not with presence of *Streptococcus* in the first infection or with detection of *S. pneumoniae* antibodies at 12 months of age. No statistically significant associations were observed for future chronic wheeze when aggregating the 7-week healthy *Streptococcus, Haemophilus* and *Moraxella* MPGs into a single predictor (with or without atopy by two years). Overall, infants who became atopic by age 2 and developed subsequent chronic wheeze at age 5 were twice as likely to have had early *Streptococcus* colonization, febrile LRI and/or HRV-C wheezy LRI in the first year of life, compared to those that did not develop chronic wheeze (**Fig. 4B**).

**Fig. 4.**
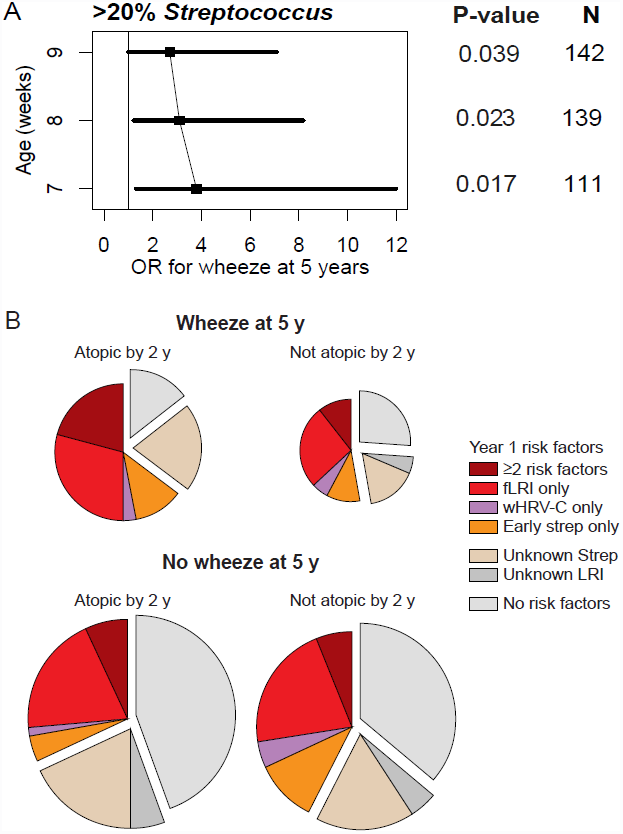
Predictors of chronic wheeze at age five. **(A)** Odds ratio (OR, filled squares) and 95% confidence intervals (black bars) for association between chronic wheeze and high (>20%) abundance of *Streptococcus* in the first healthy NP sample, estimated using logistic regression to adjust for gender, age at sample collection, maternal and paternal history of atopic disease; p-values and sample sizes are indicated to the right. Estimates were calculated separately for samples collected by 7, 8 or 9 weeks of age. Individuals who experienced an infection prior to first healthy NP sample collection were excluded from this analysis. **(B)** Distribution of microbial events during infancy that were identified as risk factors for chronic wheeze at 5 years of age, stratified according to atopic status by age 2. fLRI, febrile LRI; wHRV-C, HRV-C LRI accompanied by wheeze; Strep, >20% *Streptococcus* abundance in healthy NP sample taken in by 9 weeks old and prior to any ARI; unknown Strep, no such NP sample available, mainly due to ARI before 9 weeks of age; unknown LRI, incomplete viral/symptom profiling for LRI. Total numbers are: wheeze at 5, atopy by 2, n=34; wheeze at 5, non-atopic, 19; no wheeze at 5, atopic, 72; no wheeze at 5, non-atopic, 66. Size of piechart is proportional to the number of infants in each condition.

## Discussion

### NP microbiome composition and dynamics

Our data provide the first detailed prospective characterization of bacterial communities within the human NP microbiome during the first year of life. The NP microbiome was qualitatively simple (**Fig. 1**), dominated by six common genera - *Haemophilus*, *Streptococcus*, *Moraxella* (each more common in infection samples), *Staphylococcus*, *Alloiococcus* and *Corynebacterium* (each more common in healthy samples). This is consistent with previous studies of NP microbiome composition in children aged 12-14 months (*21*, *26*) and adults (*27*), although intra- and inter-sample diversity was greater in these older groups, likely due in part to extreme seasonal variation in those study locations. In contrast our study site in Perth, Western Australia has a very moderate climate (**Fig. S10**) and we detected only limited seasonal effects (**Fig. 2D**) that were readily adjusted for, enabling us to assess the dynamics of the NP microbiome at different stages of infancy and to examine its association with other factors.

In the CAS cohort, we found that early NP colonization typically involved *Staphylococcus* or *Corynebacterium*, which was later replaced with *Moraxella* or *Alloiococcus*. *Staphylococcus* was the dominant colonizing bacteria in the early healthy NP microbiomes, but its presence declined rapidly with age (**Fig. 1**). *S. aureus* is a common cause of neonatal sepsis (*28*) and high rates of *S. aureus* nasal colonization in the first months of life have been reported (*14*, *29*). Few studies have examined longitudinal colonization, however a recent study in African children reported *S. aureus* colonization in 42% of infants at 1 month and 12% at 12 months of age, a trend which was mirrored in maternal colonization rates (*30*) and is strikingly similar to the patterns we observed (41% at 2 months, 11% at 12 months). *Staphylococcus* and *Corynebacterium* are both common components of the human skin microbiome, and in our data they showed a comparable temporal pattern with high rates in the early NP microbiome, declining with age (**Fig. 1**). We speculate that infants tend to be colonized initially with skin-dwelling bacteria (acquired from parents and others), which is replaced over time by stable colonization with *Moraxella* or *Alloiococcus* and punctuated by transient acquisition of *Streptococcus* or *Haemophilus* that is frequently accompanied by ARI symptoms. Our data further suggests the transition to *Moraxella* is associated with exposure to other children (in the home or at day-care) and episodes of ARI, and that colonization with *Moraxella*, *Streptococcus* and *Haemophilus* are selected for by antibiotic exposure (**Fig. 2**).

*Moraxella* was the most common genus in our study population, dominating 21% of all healthy NP microbiomes and 38% of infection samples. *Moraxella* was represented in our sequence data by a single OTU (**Fig. S2**), which matches that assigned to *M. catarrhalis*, a human-restricted unencapsulated Gram negative bacterium previously associated with both commensal NP colonization and pathogenicity in the respiratory tract and inner ear (otitis media) (*31*). Increased rates of NP colonization with *M. catarrhalis* have been reported in children following pneumococcal vaccination (*32*), however the samples in our study predate introduction of this vaccine. In our cohort the presence of *Moraxella* increased with age and *Moraxella* was a particularly stable component of the NP microbiome during both health and disease (**Fig. S8**). These findings are consistent with *Moraxella*’s known ability to form biofilms (*26*, *33*), which offer protection from antibiotics (*34*) and promote co-colonization with, rather than replacement by, other common bacteria such as *S. pneumoniae* and *H. influenzae* (*35*). *Moraxella* was more abundant during the cooler months of autumn and winter, a seasonal trend consistent with recent reports that cold shock at 26oC stimulates growth, colonization and expression of virulence-associated traits in *M. cattarhalis* (*36*, *37*). These data are consistent with a Dutch microbiological culture-based study of *M. catarrhalis* carriage in young children, which found increasing prevalence of *M. catarrhalis* during the first year of life, with a strong seasonal effect (also observed in a Swedish cohort (*38*)) and a significant positive association with day-care attendance and siblings (*35*, *39*). However these studies did not examine the stability of *Moraxella* colonization within individuals over time. Importantly, our data show the presence of *Moraxella* contributes to severity of RSV respiratory infections in infants (**Fig. S13**, **Table S3**), which may be mechanistically related to RSV-*Moraxella* interactions reported in OM pathogenesis (*40*).

*Alloiococcus* was a common and stable component of healthy NP samples and demonstrated enhanced stability in healthy NP microbiomes (**Fig. S8**). *Alloiococcus* is a Gram positive bacterium with one named species, *A. otitidis,* frequently detected in the ear canals of children with OM. Little is known about its mechanisms of colonization, however the stable colonization patterns we observed may reflect a propensity to form stable biofilms, similar to *Moraxella*. The *Alloiococcus* MPG included some samples with high abundances of both *Alloiococcus* and *Corynebacterium*, indicating compatibility between these bacteria somewhat reminiscent of that between the *Moraxella* biofilm and *Streptococcus* or *Haemophilus*. Whether *Alloiococcus* is involved in OM pathology or is merely a healthy component of the microbiome is controversial (*41*); in our data, *Alloiococcus* was not associated with ARI or OM (**Fig. 1**, **Table S2**).

*Haemophilus* was almost exclusively associated with ARI symptoms and was often found in consecutive infections, suggesting it can persist in the nasopharynx for some time (**Fig. 1**, **Table 1**, **Fig. S8**). Our analyses suggest *Haemophilus* is highly transmissible between children, and selected for by antibiotic exposure (**Fig. 2**). Although our data include multiple *Haemophilus* OTUs, these epidemiological patterns are consistent with the human-adapted pathogen *H. influenzae*, which was supported by antibody data (**Fig. S1**).

### Role of the NP microbiome in asthma development

We have previously reported that within the CAS cohort, the frequency during the first year of life of severe (wheezy and/or febrile) LRI was positively associated with subsequent risk for persistent wheeze at 5 and 10 years of age (*9*, *10*,*25*), and similar findings have been reported in the US-based COAST cohort (*42*). In both these study populations the susceptible subgroup of children were those who developed early allergic sensitization (*7*-*10*, *25*, *42*, *43*), and a variety of evidence suggests that the underlying mechanism involves synergistic interactions between atopic and anti-viral inflammatory pathways triggered within the infected airway mucosa (*3*). Of note, ARIs that remain restricted to the upper respiratory tract, and infections resulting in only mild LRI without wheezing or febrile symptoms, are relatively benign with respect to asthma risk in this cohort (*9*, *10*, *25*) (**Table S3**). Hence cofactors that can enhance the spread and ensuing severity of viral-initiated ARI are potentially central to asthma pathogenesis.

Here we found that two specific classes of LRI, namely febrile LRI and HRV-C-positive wheezy LRI, were independent risk factors for later chronic wheeze, especially amongst atopics (**Table 2**, **Fig. 3**). Febrile LRI was common, occurring in one third of all children and half of atopic chronic wheezers at age 5 (**Fig. 3**–**4**). Notably, the presence of illness-associated bacteria within the NP microbiome at the time of ARI increased both the risk for progression to the lower respiratory tract and the development of fever (**Table 1**). These mechanisms likely include myriad bacterial-viral interactions, recently reviewed in (*20*). In contrast, neither specific virus groups nor bacterial MPGs were associated with wheezy vs non-wheezy LRI within the overall population (**Table 1**), and the impact of wheezy LRI on later chronic wheeze was limited to LRI with HRV-C (**Table 2**) which occurred in just 11% of children. Re-examination of wheeze symptoms during LRI stratified by atopic status identified a positive association between HRV-C and wheeze during LRI amongst those children who later became sensitized (by 2 years of age), consistent with observations that HRV-C can induce wheeze in high-risk individuals (*44*). However, the presence of wheezy LRI alone, without further stratification by HRV-C detection and later atopic status, was not a significant predictor of later chronic wheeze, while febrile LRI per se was a significant predictor of chronic wheeze at 5 and 10 years, both amongst atopics and across the whole cohort (**Fig. 4**, **Table 2**).

Early asymptomatic *Streptococcus* colonization at ~2 months of age, which occurred in 14% of children tested at that time, was significantly associated with chronic wheeze at 5 years of age (**Fig. 4**, **Table 2**). Early *Streptococcus* colonization was not associated with incidence of infections with *Streptococcus* MPG, nor with detection of *S. pneumoniae*-specific antibodies at 12 months, suggesting that the mechanism is independent of innate immune response to *Streptococcus*. Rather, early *Streptococcus* colonization was associated with younger age of first LRI, and the level of subsequent asthma risk appears inversely related to age at initial *Streptococcus* colonization (**Fig. 4**, **Fig. S14**). Collectively, these observations suggest that developing airway tissues may be maximally susceptible to the long-term effects of *Streptococcus*-mediated or LRI-mediated damage during the early postnatal period, when lung growth rates are most rapid.

To our knowledge, the only other study reporting the impact of early life NP colonization on wheeze/asthma at pre-school age is the 2007 report on the Copenhagen Prospective Study on Asthma in Childhood (n=321, children born to mothers with asthma), in which a higher rate of wheeze at five years of age was detected amongst children colonized with *S. pneumoniae, M. catarrhalis* and/or *H. influenzae* at 1 month post birth (in culture-confirmed colonized, asthma prevalence was 33%, not colonized, 10%, OR 4.57) (*14*). In that study, associations were not reported individually for the three species and colonization was measured as positive or negative by microbiological culture. In our cohort, wheeze at 5 years was not associated with early colonization with *Haemophilus* or *Moraxella*, however *Haemophilus* was very rare in healthy NP samples and *Moraxella* colonization was established later during infancy, which may be related to the warmer climate in Perth; these associations may differ in populations where asymptomatic carriage of these organisms is higher.

### Conclusions and implications

These findings collectively suggest that bacterial pathogens present in the NP microbiome at the time of upper respiratory viral infections during infancy are significant determinants of risk for the spread of infection to the lower airways and for the resultant expression of inflammatory symptoms marked by fever, and further they may contribute directly and indirectly to the ensuing risk for development of persistent asthma which itself is linked to these prior infectious episodes. Prevention of RSV or HRV infections in high-risk children, using immunoprophylaxis or vaccines, has been proposed as a mechanism for preventing the development and/or exacerbation of childhood asthma (*12*, *42*, *45*). Our data suggests that in the absence of effective anti-viral therapies, targeting pathogenic bacteria present within the NP microbiome in this age group could represent an alternative approach towards the same goal. The pneumococcal vaccine, currently recommended from 2 months of age, could play a role; however the niche created by eliminating vaccine-targeted *S. pneumoniae* serotypes can be readily filled by other serotypes (*46*) and other bacteria such as *H. influenzae* (*26*) and *M. catarrhalis* (*32*). Manipulation of the microbiome by antibiotics may appear attractive, however the association between antibiotic use in early childhood and subsequent asthma is controversial. In the CAS cohort (*22*) and others (*47*), this association has been proposed to arise through confounding and reverse causation, whereby genetic or clinical factors increase the likelihood of both antibiotic prescription (e.g. via genetic susceptibility to viral infection (*47*)) and asthma development; others have proposed that antibiotic-induced disruption of the gut microbiota could explain the link (*48*). Our analyses provide a potential causal pathway linking antibiotics to later asthma, whereby antibiotic use in infants selects for illness-associated bacteria in the NP microbiome, leading to increased risk of febrile LRI and later asthma development. Genetic analysis of the infants in our cohort could potentially help to unravel this in the future. Importantly, further studies are required to prospectively assess the impact of antibiotic use on NP bacterial colonization during health and infection, and the subsequent development of ARI, atopy and wheeze.

## Acknowledgments

This study was supported by the NHMRC of Australia (Project Grant #1049539; Fellowships #1061409 (KEH) and #1061435 (MI, co-funded with the Australian Heart Foundation) and the Victorian Life Sciences Computation Initiative (VLSCI) (#VR0082). SLJ is supported in part by a Chair from Asthma UK (CH11SJ) and by Medical Research Council Centre grant G1000758. JEG, YAB and KG are supported by NIH grants U19 AI104317 and P01 HL070831.

